# Oxytocin receptor and HER2 interactions in breast cancer

**DOI:** 10.1101/2025.03.09.642136

**Authors:** Huiping Liu, Liying Liu, Andreas Möller, Markus Muttenthaler

## Abstract

Breast cancer affects women globally, with HER2 being one of the most aggressive subtypes. Despite advances in HER2-targeted therapies, many patients fail to respond to such treatments. In search of alternative therapeutic targets, we investigated the role of the oxytocin (OT) and oxytocin receptor (OTR) signalling system in HER2^+^ breast cancer. Survival analysis of HER2 subtype breast cancer patients revealed high OTR expression correlating with significantly improved relapse-free survival. OTR overexpression or knockdown in HER2^+^ SK-BR-3 cells decreased or increased cell viability, respectively. HER2 was downregulated in OTR-overexpressing cells, which could contribute to the decreased Pertuzumab efficacy. Importantly, HER2 was internalised upon OT treatment and interacted with OTR to form OTR-HER2 complexes. OT treatment furthermore induced a transient inhibition of HER2 phosphorylation without affecting the total HER2 protein levels. In addition, OT inhibited ERK1/2 phosphorylation but enhanced Akt phosphorylation, linked to increased cell viability. In summary, this work describes a novel HER2-OTR interaction and mechanism, expanding our understanding of HER2 cancer biology that might lead to new prognostic and therapeutic opportunities centred around the OT/OTR signalling system for patients suffering from HER2^+^ breast cancer.

## Introduction

Breast cancer comprises several molecularly and biologically heterogeneous disease subtypes that have different clinical prognoses. The HER2 (human epidermal growth factor receptor 2) subtype is characterised by HER2 overexpression in these cancer cells, accounting for ∼20% of all breast tumours^1^. HER2 is a member of the human epidermal growth factor receptor (EGFR/ErbB/HER) family, which consists of four plasma membrane-bound receptor tyrosine kinases (RTK), including EGFR (ErbB-1), HER2/neu (ErbB-2), HER3 (ErbB-3), and HER4 (ErbB-4)^2^. HER2 is an orphan receptor with no known natural ligands. However, HER2’s extracellular domain contains a constitutively active conformation that mimics the activated ligand-bound state of EGFR^3^. HER2 homodimerization or heterodimerization with other receptors of the HER family results in activation of mitogenic signalling, and HER2 overexpression leads to constitutive activation of the HER2 signalling pathways^4^. HER2 breast tumours are aggressive, and HER2 breast cancer patients have had poor prognosis and clinical outcomes before HER2-targeted therapies were introduced ^5^. Despite an improvement in patient survival due to HER2-targeted treatments, a considerable number of patients relapse or develop progressive disease due to either inherent or acquired drug resistance^6^, highlighting the need of alternative strategies and targets.

The oxytocin receptor (OTR) is a class-A G protein-coupled receptor (GPCR) that is highly expressed in the mammary glands, where it mediates the effects of oxytocin (OT) during lactation. Emerging evidence supports a functional role of OT/OTR also in breast cancer development and progression,^7^ including in the HER2^+^ subtype.^8, 9^ OT/OTR’s function in HER2^+^ breast cancer remains, however, poorly understood. GPCRs are known to crosstalk with the HER/EGFR family, and HER2 has been reported to interact with GPCRs such as the β2-adrenergic receptor (β2AR)^10^ and the cannabinoid receptor 2 (CB2R),^11^ resulting in unique signalling with (patho)physiological implications. OTR also forms heterodimers with other receptors, for example, with the β2AR in myometrial cells,^12^ with the dopamine 2 receptor (D2R) in the dorsal and ventral striatum and in the nucleus,^13^ or with the vasopressin receptors during biosynthesis.^14^

In this study, we investigated OT/OTR in the HER2 breast cancer subtype, revealing new mechanistic insights into the OT/OTR signalling system on disease pathogenesis and progression and providing new opportunities for disease management.

## Materials and Methods

### OT/OTR mRNA level and breast cancer patient survival

The Kaplan–Meier Plotter-Breast Cancer (KM-Plotter, www.kmplot.com) online platform was used to assess the prognostic values of OT/OTR mRNA (mRNA, gene Chip) across different intrinsic breast cancer subtypes (basal, luminal A, luminal B and HER2).^15, 16^

### Cell culture and cell transfection

Human breast cancer cell line SK-BR-3 (HER2^+^) was routinely cultured in RPMI 1640 (Life Technologies Limited, Grand Island, NY, USA) supplemented with 10% fetal bovine serum (FBS, GE Healthcare Life Sciences, Mordialloc, VIC, AUS) at 37°C in a humid atmosphere of 5% CO_2_, routinely confirmed negative for mycoplasma and bacteria contamination. The cell line was a gift from Prof Andreas Möller (QIMR Berghofer Medical Research Institute). All experiments were performed with mycoplasma-free cells.

OTR overexpression was achieved in SK-BR-3 cells by transient transfection of expression vectors with or without the OTR sequence (pCMV6-OTR and pCMV6-vector obtained from Origene, Rockville, MD, USA) with FuGENE HD transfection reagent (Promega, Madison, WI, USA) into human breast cancer cells. Transient genetic knock-down was performed by selective siRNA SMARTpool (which is a combination of four SMART selection-designed siRNAs into a single pool) transfection with DharmaFECT 1 transfection reagent (Dharmacon, Horizon Discovery, Lafayette, CO, USA). The control (non-targeted) siRNA was also purchased from Dharmacon (place) as a SMARTpool.

### MTT assay

Cell proliferation was assessed using the 3-(4,5-dimethyl-2-thiazolyl)-2,5-diphenyl-2H-tetrazolium bromide (MTT) assay, a colourimetric assay for assessing viable cell metabolic activity. Cells were collected and seeded in 96-well plates (5×10^3^ cells per well). OT was synthesised in our lab with a purity of >98%, measured by high-performance liquid chromatography (HPLC). Pertuzumab is a HER2-targeted monoclonal antibody purchased from Roche. The treatments were implemented after cell adherence by replacing the medium in the plate with fresh phenol red-free medium supplemented with 5% charcoal-stripped FBS containing different treatments. Controls were treated with an equal volume of vehicle (H_2_O) without compounds. MTT (5 mg/mL in PBS, 20 μL/well, Sigma-Aldrich, Bayswater, VIC, Australia) was added into the wells after 72 h, and the plates were incubated at 37°C for 4 h. After centrifugation of the plates, the supernatants were removed and DMSO (150 μL/well) was added. Light absorbance of the solution was then measured at λ = 570 nm on an INFINITE M1000 PRO plate reader (Tecan Austria GmbH, Grödig, Austria).

### Western blots

Cells were seeded into 48-well plates (5×10^4^ cells per well) and serum-starved overnight before stimulation. Cells were treated with 1 μM peptides for 5, 15, 30, 60, 120, 180, and 240 min, washed with ice-cold PBS, and then lysed with 1× laemmli buffer containing 50 mM DTT (Sigma-Aldrich, North Ryde BC, NSW, AUS). Protein concentration was determined using a Pierce™ BCA Protein Assay Kit (Thermo Fisher, Rockford, IL, USA). The same amount of sample (∼20 μg) was loaded and separated by SDS-PAGE gel electrophoresis and transferred onto Polyvinylidene Fluoride (PVDF) membrane (Merk Millipore Ltd., Ireland). The membrane was probed with the following primary antibodies: mouse-antibody for DYKDDDDK tag (FLAG) (MA191878, Invitrogen, Rockford, IL, USA), rabbit anti-HER2 (2165; Cell Signaling Technology, Beverly, MA, USA), rabbit anti-p-HER2 Tyr^1248^ (2274; Cell Signaling Technology), mouse anti-p44/42 MAPK (Erk1/2) (4696; Cell Signaling Technology), rabbit anti-Phospho-p44/42 MAPK (Erk1/2) (Thr^202^/Tyr^204^) (4370; Cell Signaling Technology), rabbit-anti-Akt (4691; Cell Signaling Technology), rabbit anti-Phospho-Akt (4060; Cell Signaling Technology), mouse anti-tubulin (M30109S; Abmart), and mouse anti-GAPDH (sc-47724; Santa Cruz). GAPDH or tubulin was used as loading controls. Membranes were then incubated with Donkey anti-mouse IRDye680 or Donkey anti-rabbit IRDye800 labelled secondary antibody (LI-COR, Lincoln, NE, USA), imaged and analysed using ODYSSEY Infrared Imaging System (LI-COR, Lincoln, NE, USA).

### Co-immunoprecipitation assay

Co-immunoprecipitation was conducted using a Pierce™ Classic Magnetic IP/Co-IP Kit (Thermo Fisher Scientific, Scoresby, VIC, Australia) according to the manufacturer’s instructions. Briefly, cells were lysed with a lysis/wash buffer provided with the kit, and the protein concentration was tested using the Pierce™ BCA Protein Assay Kit (Thermo Fisher Scientific, Scoresby, VIC, Australia). A total protein amount of 500 μg (input) was incubated overnight at 4°C with 5 μg of IP antibody (anti-FLAG antibody specifically targeting the OTR-FLAG and the mouse IgG as negative control) per sample to prepare the immune complex. For each sample, 25 μL (0.25 mg) of Pierce Protein A/G Magnetic Beads were added for immunoprecipitation at 25°C for 1 h. After separating the beads using a magnetic stand, the immunoprecipitated proteins were eluted with a Lane Marker Sample Buffer containing 50 mM DTT. The resulting samples were separated by SDS-PAGE electrophoresis and immunodetected using HER2 and FLAG-specific antibodies by western blot.

### Immunofluorescence analysis by confocal microscopy

SK-BR-3 (wild-type) cells were grown on coverslips in 24-well plates (2–5×10^4^ cells per well). Cells were treated with 1 μM OT for 5 min or 24 h. After treatment, cells were washed with ice-cold PBS and fixed with 4% paraformaldehyde for 20 min. Cell samples were permeabilised with 0.1% Triton X100 for 5 min, washed three times with PBS and blocked with 0.5% BSA/PBS for 30 min. Primary rabbit anti-HER2 antibody (2165; Cell Signaling Technology) was prepared in 0.5% BSA/PBS and incubated for at least 30 min. The cells were washed three times with PBS and incubated with Alexa Fluor 488-conjugated goat anti-rabbit secondary antibody (Invitrogen, Mulgrave, VIC, Australia) for 30 min at 25°C. After washing, a drop of Prolong Gold antifade reagent with DAPI (Invitrogen, Mulgrave, VIC, Australia) was added onto a clean glass slide, and the dry coverslip was placed cells-side down onto mounting media. The slide was then sealed with a sealant. All images were obtained using a Fluoro 2 – Zeiss Apotome 2.0 confocal microscope and settings were adjusted to allow for the detection of the cell membrane structure. As a result, fluorescent intensities could not be used for quantitation.

### Statistical analysis

Kaplan–Meier survival curves were statistically compared using the log-rank test. Log-rank p-value and hazard ratio (HR) with 95% confidence intervals were calculated and displayed on the plot figures. Pearson correlation analysis was applied to determine the association of the mRNA expression (RNAseq) of OTR and HER2 in the 57 breast cancer cell lines from the Cancer Cell Line Encyclopedia (CCLE, https://portals.broadinstitute.org/ccle). For the comparison of multiple groups, one-way ANOVA was used, followed by Dunnett’s multiple comparison test. Student’s t test was applied when comparing two groups. Pearson’s correlation analysis was used to determine correlations. GraphPad Prism (version 8.4.2, San Diego, CA, USA) was used for all data analysis and figure plotting. Data are presented as mean ± SEM of at least three independent assays unless otherwise specified. The significance level was considered below 0.05 in all experiments.

## Results

### Opposing roles of OT and OTR in HER2 breast cancer subtype patient survival and cell proliferation

We investigated the correlations of OT and OTR expression in predicting relapse-free survival (RFS) of different breast cancer subtypes using the KM-Plotter. Opposing prognostic values for OT and OTR were observed with patients having the HER2 subtype, with high OTR levels correlating with better RFS probability (HR 0.64 [0.43-0.95], p = 0.027, n = 251), while high OT levels correlated with lower RFS probability (HR 1.42 [0.94-2.15], p = 0.092, n = 251, Fig. 1A). The upper quartile survival for HER2 breast cancer patients with high or low OTR expression was 22.6 or 16.0 months, respectively, while with high or low OT expression, it was 14.0 or 20.4 months, respectively (Fig. 1E).

**Figure 1.**
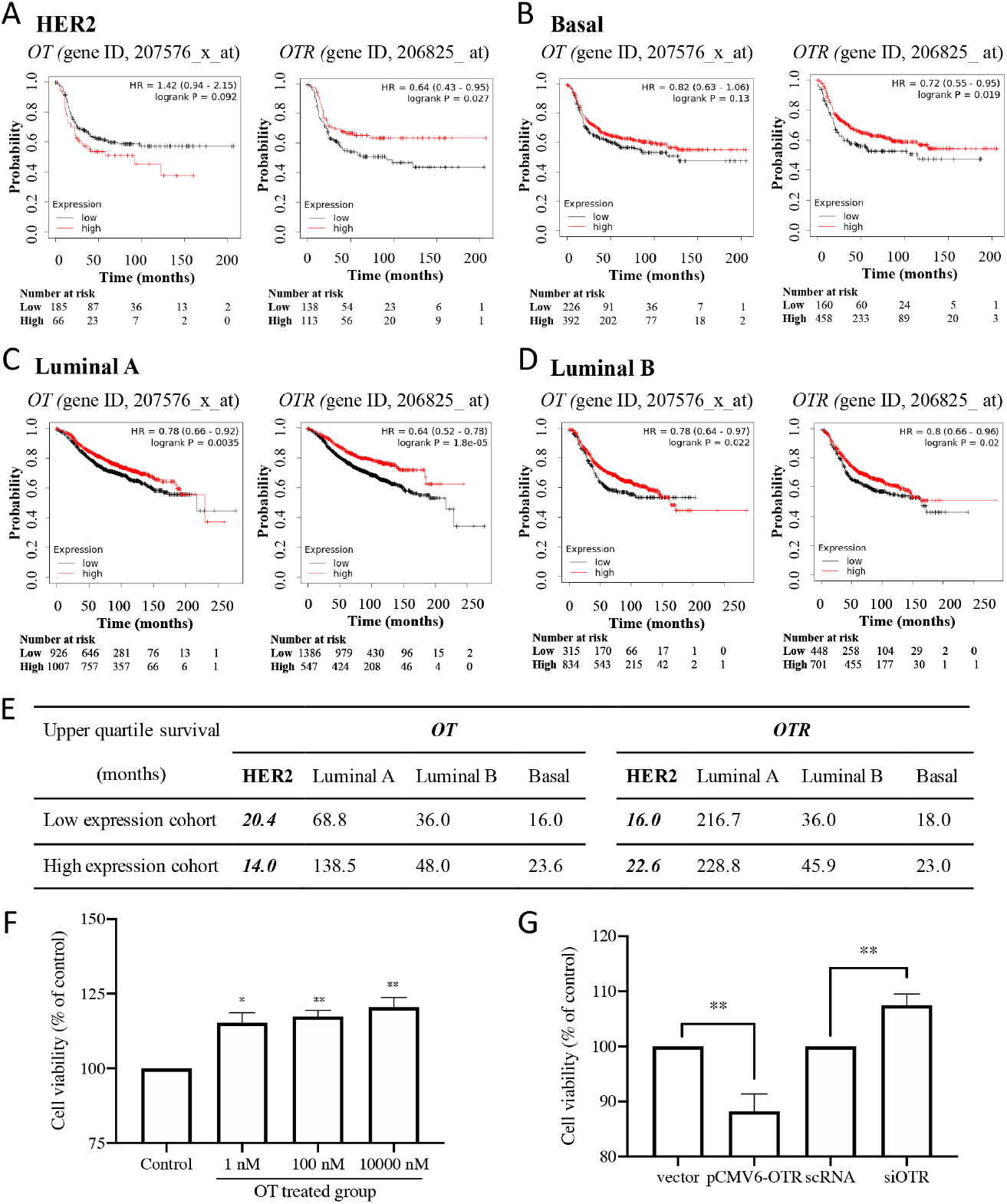
Roles of oxytocin and oxytocin receptor in breast cancer patient survival based on Kaplan Meier-plotter and HER2 subtype cell proliferationn. **A-D**. Relapse-free survival (RFS) probability curves of HER2/basal/luminal A/luminal B subtype breast cancer patients with high or low OT/OTR expression. The auto-selected best cut-off button was chosen so that patients were stratified into high- and low-level groups according to the best cut-off, which was the OT/OTR expression value that had the best-performing threshold in survival analysis. The ‘Number at risk’ (shown below the plot to the corresponding time point) indicates the number of patients that are not censored or dead at the given time point (0, 50, 100, 150, 200, and 250 months). HR, hazard ratio; OT, gene ID, 207576_x_at; OTR, gene ID, 206825_at. **E**. Upper quartile survival summary of patients with low or high OT and OTR levels in different breast cancer subtypes. ^#^, the upper quartile survival was measured as months and rounded off from data obtained from KM-Plotter data analysis. **F**. Effects of OT treatment on *in vitro* cell viability of breast cancer cell lines using the MTT assay. Cells were incubated with the indicated doses of OT for 72 h, and cell viability was assessed using the MTT assay after the treatment. Statistical analysis used One-way ANOVA analysis followed by Tukey’s multiple comparisons test. **G**. Effects of OTR expression modulation on SK-BR-3 cell growth. Cells were seeded in a 96-well plate and transfected with the pCMV6-OTR plasmid for OTR overexpression or siRNA SMARTpool for OTR knockdown (siOTR). After 24 h, the transfected cells were allowed to continue in culture for 48 h before adding MTT. Statistical analysis was using the Student’s t-test. **, p < 0.01 vs control (vector group was the control for pCMV-OTR group, and scRNA (scrambled non-targeting RNA) group was the control for siOTR group). *, p < 0.05; **, p < 0.01; ***, p < 0.001 vs control; n ≥ 3.

We pursued these findings on the cellular level using the HER2 breast cancer cell line SK-BR-3, which expresses both OT and OTR mRNA.^17^ OT treatment (1 nM, 100 nM, 10 μM) increased cell viability to 115 ± 3% (p < 0.05), 117 ± 2% (p < 0.001) and 120 ± 3% (p < 0.01) after 72 h compared to the control group with vehicle treatment (100%), respectively (Fig. 1F). By contrast, OTR overexpression reduced cell viability by 10% (p < 0.01) compared to the control group transfected with the vector plasmid (Fig. 1G This OTR-specific effect was confirmed by OTR knockdown experiments, which enhanced cell viability by 14% (p < 0.05) compared to the control group transfected with scrambled non-targeting RNA (Fig. 1G). These results aligned with the opposing prognostic values of OT and OTR observed in HER2 subtype breast cancer patients.

### OTR overexpression decreases HER2 expression in HER2+ breast cancer cells

We then assessed the OTR and HER2 mRNA profiles of 57 breast cancer cell lines from the Cancer Cell Line Encyclopedia database. An inverse association was observed between OTR and HER2 levels, with cell lines with high HER2 expression tending to have low OTR expression (Fig. 2A, Table S1). This indicated that OTR overexpression might reduce SK-BR-3 cell viability by downregulating HER2 expression. Indeed, HER2 expression was lower in SK-BR-3 cells transfected with pCMV6-OTR than in control cells on mRNA and protein levels (Fig. 2B&C, p < 0.05), supporting a connection, if not a functional interaction, between OTR and HER2. We also detected the effect of OT and OTR expression on the cell viability and cellular response to HER2-targeted antibody Pertuzumab, which binds to an extracellular domain of the HER2 receptor and inhibits heterodimerization.^18^ While OT did not significantly affect the efficacy of Pertuzumab for SK-BR-3 cell proliferation regardless of OTR expression level (Fig. 2 D&E), OTR overexpression significantly decreased Pertuzumab efficacy when used alone (Fig. 2F, p < 0.05), which could be partly attributed to the decreased levels of HER2 on the cell membrane.

**Figure 2.**
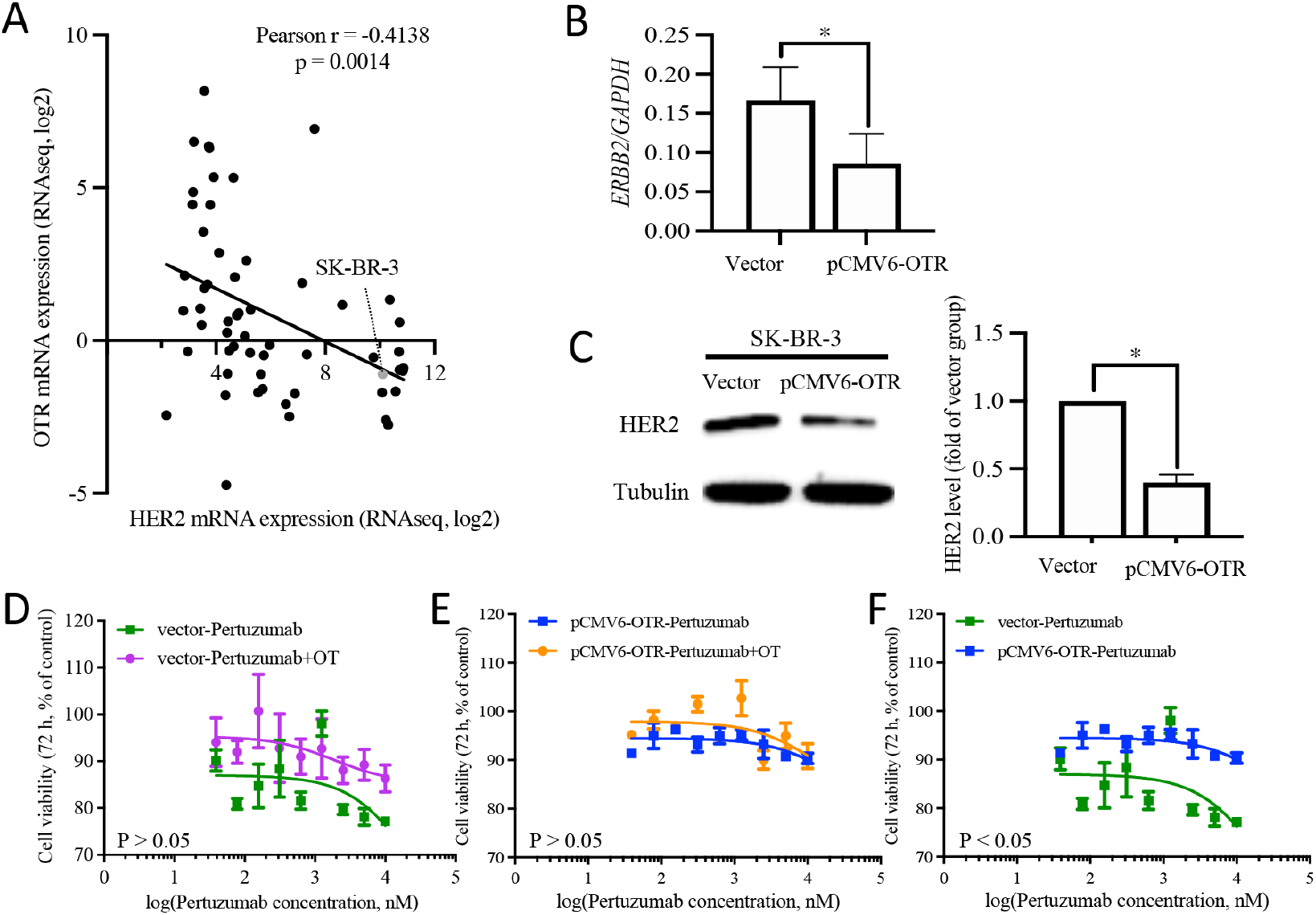
Oxytocin receptor and HER2 expression in breast cancer cells. **A**. Correlation between OTR and HER2 mRNA levels in 57 breast cancer cell lines from the Cancer Cell Line Encyclopedia as analysed by Pearson correlation analysis. Each dot indicates one cell line, with the SK-BR-3 cell line labelled grey. **B & C**. HER2 expression on mRNA and protein levels in OTR-overexpressing SK-BR-3 cells. Cells were transfected with the pCMV6 plasmid vector or the pCMV6-OTR plasmid for the control group or OTR overexpression, respectively. HER2 mRNA and protein levels were analysed by qPCR and western blot assay 24 h after transfection (GAPDH was used as an internal control for gene expression, and tubulin was used as an internal control for protein expression, n = 3). **D - F**. Effects of Pertuzumab on SK-BR-3 cell proliferation with or without OT. The Student’s t-test was used for statistical analysis *, p < 0.05; **, p < 0.01; ****, p < 0.0001.

### OT-induced HER2 phosphorylation and trafficking

GPCR stimulation can lead to activation and intracellular phosphorylation of RTKs (such as EGFR), which triggers cellular functions, including cell growth, differentiation, motility, or death.^19-22^ No HER2 activation was observed upon OT treatment, but HER2 phosphorylation was reduced by ∼60% upon 5 min of OT treatment, which gradually returned to baseline after 60 min (Fig. 3A). Since total HER2 did not decrease upon OT treatment, we hypothesised that the reduction in HER2 phosphorylation could be related to a reduction of cell surface HER2. We therefore investigated HER2 localisation upon OT treatment by confocal microscopy. Interestingly, OT induced HER2 internalisation in SK-BR-3 cells within 5 min (Fig. 3B). After 24 h, HER2 was detected at the cell membrane again, with both results suggesting a transient effect of OT on HER2 function (Fig. 3B).

**Figure 3.**
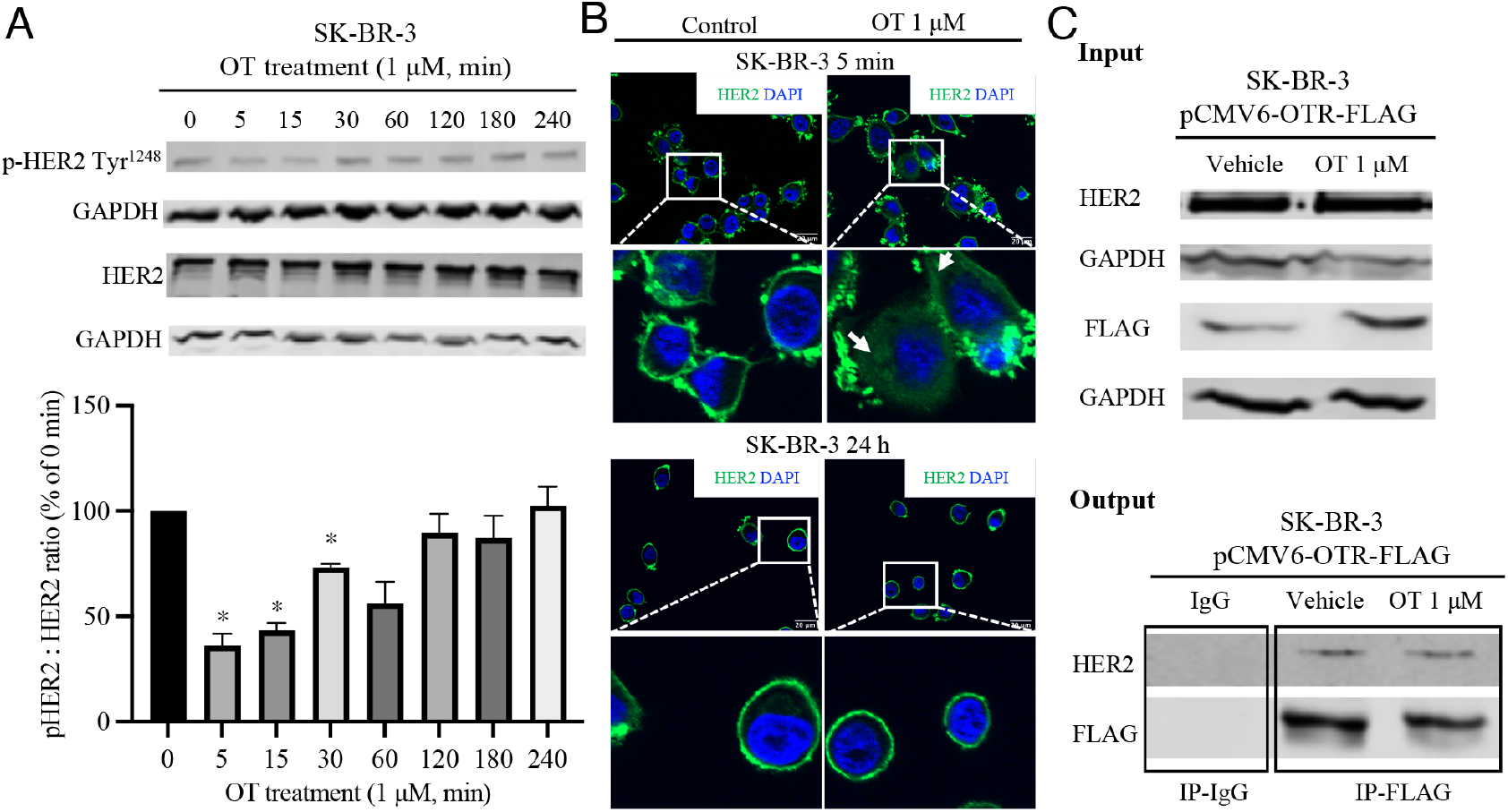
Effect of oxytocin on HER2 phosphorylation and protein trafficking in SK-BR-3 cells. **A**. Time course of HER2 phosphorylation in response to OT (1 μM) treatment detected by western blot. The bar graph represents the quantification of the phosphorylation indicated by the pHER2:HER2 ratio. One-way ANOVA analysis followed by Tukey’s multiple comparisons test was used for statistical analysis. *, p < 0.05 vs. 0 min group without OT treatment; n ≥ 3. **B**. Confocal microscope images of immunofluorescence for HER2 (green) with DAPI nucleus staining (blue) in SK-BR-3 cells treated with OT (1 μM) for 5 min and 24 h. HER2 was visualised with a primary rabbit antibody that specifically binds to the HER2 protein, followed by an Alexa Fluor 488-conjugated goat anti-rabbit secondary antibody using a Fluoro 2 – Zeiss Apotome 2.0 confocal microscope. The magnification for the top panel images is 630× (scale bar 20 μm), and the bottom panel images represent enlargements of the boxed areas from the top panel. Control, cells without OT treatment. **C**. Co-immunoprecipitation (Co-IP) demonstrated interaction of OTR with HER2. Total protein extracts from the cell lysis (input) were immunoprecipitated with anti-FLAG antibody (targeting the OTR-FLAG) and the mouse IgG (negative control) and resolved on SDS-PAGE gel. Protein interaction was immunodetected using HER2 and FLAG antibodies. Heterodimer of HER2 and OTR-FLAG was observed (output).

We next wondered whether OT impacts the functional crosstalk between HER2 and GPCRs, which relies on physical interactions between the receptors, such as the heterodimerization of HER2 with β2AR or CB2R.^10, 11^ We hypothesised that HER2 might form a heterodimer with OTR and that this complex is internalised upon OT stimulation. Hence, we investigated this potential HER2-OTR complex formation *via* co-immunoprecipitation. SK-BR-3 cells were transfected with a FLAG-tagged OTR plasmid (pCMV6-OTR), and a FLAG antibody was used to pull down the protein complex containing OTR-FLAG. The pulled-down protein complex contained significant amounts of HER2, confirming an OTR/HER2 complex in breast cancer cells (Fig. 3C). No difference was observed between vehicle and OT treatment, suggesting that the OTR-HER2 complex was pre-formed independent of OT stimulation and that the OTR-HER2 complex stayed intact during and after internalisation.

### OT effects on ERK1/2 and Akt phosphorylation

Both HER2 and OTR can activate the Ras-Raf-ERK1/2 and PI3K-Akt pathways that play important roles in cell proliferation and survival (Fig. 4A). We therefore investigated the effect of OT treatment on ERK1/2 and Akt pathways. OT treatment resulted in ERK1/2 phosphorylation at 5 min, followed by a significant (p < 0.01) inhibition of ERK1/2 phosphorylation after 30 min (Fig. 4B). Intriguingly, OT only inhibited ERK1/2 phosphorylation in SK-BR-3 cells but not in luminal B BT474, basal triple-negative MDA-MB-231 and luminal A MCF-7 cell lines, suggesting breast cancer subtype specificity (Fig. S1) potentially relying on the presence of the HER2/OTR complex. OTR activation by OT either induces transient or sustained ERK1/2 activation,^23^ and we are not aware of any study reporting ERK1/2 inhibition by OT. The slight ERK1/2 activation at 5 min (p > 0.05) likely relates to OT-OTR activation,^23-25^ while the later reduction in ERK1/2 phosphorylation could be due to the above-demonstrated reduced HER2 phosphorylation caused by OT-induced OTR-HER2 internalisation. In addition, OT treatment increased Akt phosphorylation at 5–15 min (p < 0.01), which returned to baseline at 30 min (Fig. 3C). These observations align with the pro-oncogenic role of OT (ERK1/2 and Akt activation) and with the protective role of OTR (reduction in ERK1/2 phosphorylation due to HER2-OTR internalisation).

**Figure 4.**
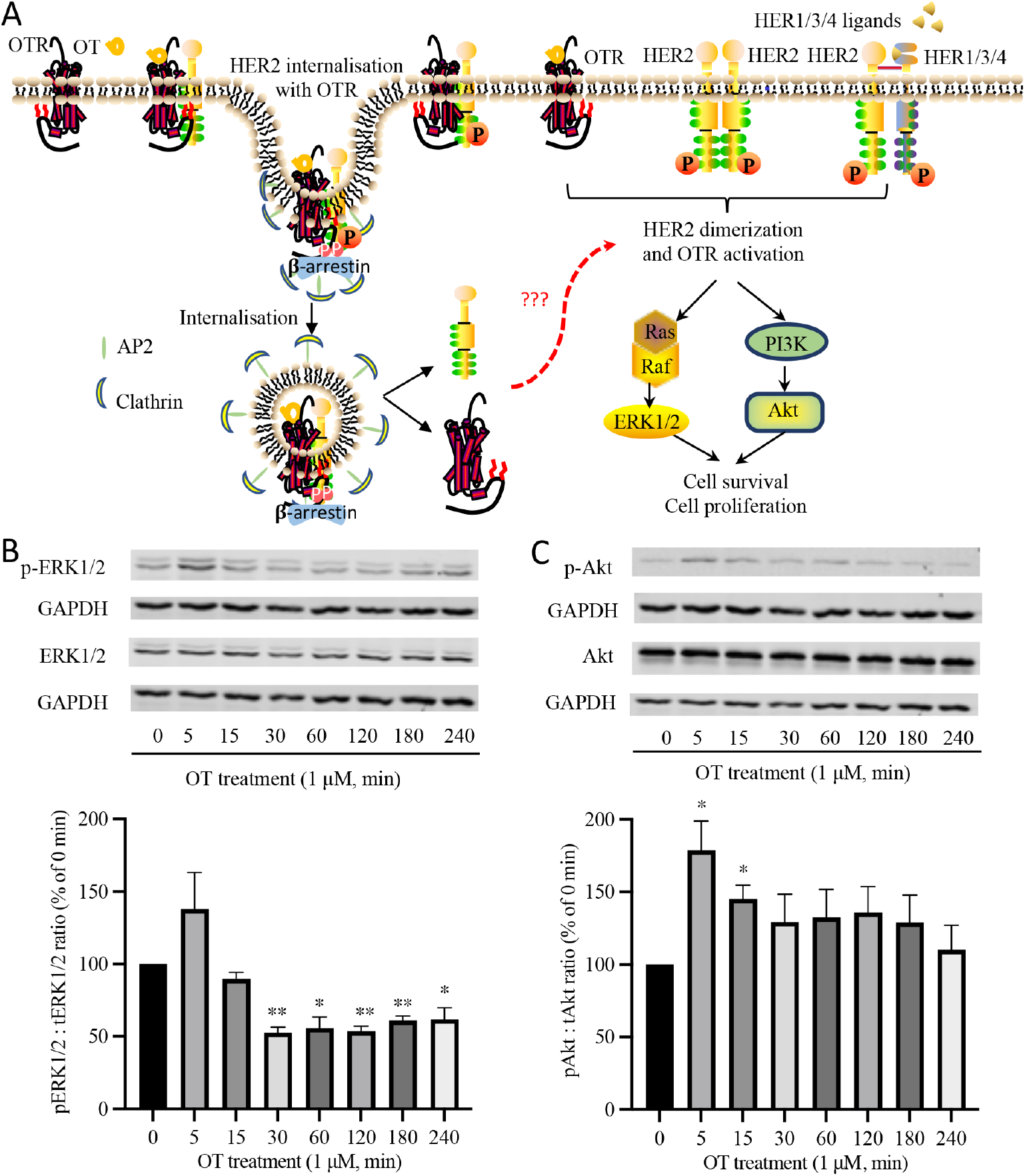
OT effects on ERK1/2 and Akt phosphorylation in HER2^+^ SK-BR-3 cells. **A**. Proposed model of HER2-OTR internalisation. OTR activation induces ERK1/2 and Akt phosphorylation. HER2-mediated signalling is either initiated by homodimerization when HER2 is expressed at high levels or by ligand-induced heterodimerization with other HER family receptors (HER1/HER3/HER4). Receptor dimerization causes transphosphorylation of tyrosine residues within the intracellular domains of the HER family receptors. Phosphorylation of HER2 reveals its binding sites for adaptor or effector proteins such as Ras/Raf and PI3K, which further activate ERK and Akt pathways, promoting cell survival and proliferation. **B & C**. Representative western blot images of total and phosphorylated ERK1/2 and Akt induced by OT treatment (1 μM). Column figures represent the quantification of the phosphorylation indicated by pERK1/2:ERK1/2 or pAkt:Akt ratio compared with 0 min group (n = 3 independent experiments). One-way ANOVA analysis followed by Tukey’s multiple comparisons test was used for statistical analysis. *, p < 0.05, **, p < 0.001 vs 0 min group (without OT treatment).

## Discussion

The OT/OTR signalling system has been investigated in several studies, supporting a mediating role during breast cancer development and progression.^7, 26^ The current consensus is that high OT and OTR levels display overall protective effects against breast cancer; however, this has yet to be investigated in a subtype-specific context where OT/OTR effects might differ. For example, breastfeeding, during which the mammary gland OTR expression and OT plasma level are increased, reduces the risk of developing breast cancer,^27^ and animal studies targeting OTR for breast tumour reduction have also provided promising results.^7^ In basal, luminal A, and luminal B subtypes, high OT and OTR expression also correlate with improved patient survival (Fig. 1), aligning with the consensus of OT/OTR having protective effects. By contrast, in TNBC, high OTR expression has been linked to increased cell migration and reduced patient survival.^28^ The data of this study indicate that the OT/OTR signalling system functions differently in the HER2 subtype, considering the opposing prognostic roles of OT and OTR (high OT– reduced survival; high OTR– improved survival; Fig. 1A) along with the OT and OTR knockdown/overexpression cell viability experiments (Fig. 1 F&G), further supporting the notion that OT/OTR should generally be investigated in a subtype-specific context.

The inverse correlation of OTR and HER2 gene expression in multiple breast cancer cell lines (Fig. 2A, Table S1) and HER2 downregulation in OTR-overexpressing SK-BR-3 cells (Fig. 2B&C) add a new mechanistic angle to OTR’s role in regulating the HER2^+^ breast cancer proliferation. However, more research into the role of OTR-mediated HER2 signalling changes is required, considering the conflicting data on how OTR and HER2 can affect each other’s expression. For instance, OTR mRNA levels are higher in luminal breast tumours with lower HER2 IHC scores,^8^ while OTR is lower in TNBC and ER^+^ breast cancers compared to HER2^+^ breast cancer.^29^ OTR overexpression also induces the formation of mammary tumours with HER2 upregulation in mice,^9^ which is contrary to its assumed protective role against breast cancer in humans. On the other hand, OTR overexpression decreased HER2 levels in breast cancer cells, which could, in turn, interfere with the efficacy of HER2-targeted therapy (Fig. 2F). Further research into the crosstalk between the OTR and HER2 pathways is warranted and could yield better intervention strategies for individuals diagnosed with different levels of HER2.

HER2 is known to interact with other membrane receptors, forming an essential determinant of cell responses to various stimuli, including HER2 transactivation in response to GPCR ligand stimulation.^30^ The mechanisms of such GPCR and HER2 interactions remain, however, underexplored. Two well-characterised examples of such crosstalk are the HER2-β2AR complex in the heart, essential for cardiac homeostasis,^10^ and the HER2-CB2R heterodimer in breast cancer, a potential new marker and drug target for patients with the HER2^+^ subtype.^11^ Here, we revealed that OT induced HER2 internalisation in SK-BR-3 cells, even though HER2 typically resists internalisation and degradation.^31-33^ By contrast, OTR is efficiently internalised upon OT binding (receptor internalisation t_1/2_ = 2.3 min)^34^ and recycled *via* a short cycle (cell surface receptor binding returned to baseline around 45 min),^35^ supporting our hypothesis that HER2 internalisation was triggered *via* OTR. The OTR-HER2 complex formation (independent of OT) was confirmed by co-immunoprecipitation, providing the first evidence of a direct OTR-HER2 interaction (Fig. 3C). This is the first study demonstrating that HER2 can form a complex with OTR, independent of OT.

HER2 internalisation could explain the transient inhibition of HER2 phosphorylation upon OT treatment in SK-BR-3 cells (Fig. 3A) since HER2 usually remains on the cell surface and signals for prolonged periods after activation.^36^ These findings support a direct interaction between HER2 and OTR at the molecular level that should affect downstream signalling and breast cancer growth. The ERK1/2 and Akt pathways are two important cancer downstream pathways that were then investigated in the context of OTR and HER2 signal transduction controlling cell proliferation (Fig. 4), indicating that the Akt pathway plays a more dominant role in HER2-mediated oncogenic signalling for promoting cell growth because the inhibition of the ERK1/2 pathway did not decrease cell proliferation, consistent with previous findings.^37^

Considering the literature and this study’s data, we propose the following model for OTR-HER2 function in HER2^+^ breast cancer: HER2 can form a complex with OTR, which internalises with OTR upon OT treatment, causing transient inhibition of HER2 phosphorylation. HER2 cycles back to the cell surface within 60 min. Due to HER2 internalisation upon OT treatment, ERK1/2 phosphorylation is inhibited, an effect accompanied by enhanced and maintained Akt activation that induces cell proliferation. Open questions remain, including whether this OTR-HER2 interaction is also observed in other HER2^+^ cell lines, how OTR and HER2 pathways are regulated after internalisation, and whether or how OTR overexpression and OT treatment could interfere with dimer formation between HER2 and the HER family receptors.

## Conclusions

Taken together, we report an unprecedented complex formation between OTR and HER2 that leads to HER2 internalisation upon OT treatment, supporting OTR as a pivotal regulator of HER2 signalling. This new OTR-HER2 interaction adds to our understanding of HER2 breast cancer biology and could provide new opportunities for prognostic marker discovery and new therapeutic strategies for personalised treatment.

## Acknowledgements

Funding supporting the studies is listed as follows: M.M. was supported by the European Research Council under the European Union’s Horizon 2020 research and innovation program (714366) and by the Australian Research Council (DP190101667). M.M. and A.M. were supported by grants from Cancer Australia and Cancer Council Queensland (1146504). A.M. was further supported by the Innovation and Technology Commission, Hong Kong SAR (PiH/048-050/22GS), the Global STEM scheme (GSP153) and the Hong Kong Jockey Club Charities Trust, and the Chinese University of Hong Kong (IDBF23MED14).

H.L. was supported by the Shandong Provincial Natural Science Foundation (ZR2023QH237) and the National Natural Science Foundation of China (82404702).

## Author contributions

M.M. and H.L. designed the experiments. H.L. and L.L. performed the experiments and collected the data. M.M. and A.M. supervised and obtained research funding for this study. M.M. and H.L. analysed and interpreted the results. M.M. and H.L. wrote the manuscript. All authors have read and agreed to the published version of the manuscript.

## Data availability

All data generated or analysed during this study are included in this published article [and its supplementary information files].

## Competing interests

The authors declare that there are no competing interests.

## Notes

### Competing Interest Statement

The authors have declared no competing interest.

## References

1. Tinoco G, Warsch S, Gluck S, Avancha K, Montero AJ. Treating breast cancer in the 21st century: emerging biological therapies. J Cancer 2013;4: 117–132.

2. Wieduwilt MJ, Moasser MM. The epidermal growth factor receptor family: biology driving targeted therapeutics. Cell Mol Life Sci 2008;65: 1566–1584.

3. Garrett TP, McKern NM, Lou M, Elleman TC, Adams TE, Lovrecz GO, Kofler M, Jorissen RN, Nice EC, Burgess AW, Ward CW. The crystal structure of a truncated ErbB2 ectodomain reveals an active conformation, poised to interact with other ErbB receptors. Mol Cell 2003;11: 495–505.

4. Olayioye MA. Update on HER-2 as a target for cancer therapy: intracellular signaling pathways of ErbB2/HER-2 and family members. Breast Cancer Res 2001;3: 385–389.

5. Iqbal N, Iqbal N. Human Epidermal Growth Factor Receptor 2 (HER2) in Cancers: Overexpression and Therapeutic Implications. Mol Biol Int 2014;2014: 852748.

6. Rexer BN, Arteaga CL. Intrinsic and acquired resistance to HER2-targeted therapies in HER2 gene-amplified breast cancer: mechanisms and clinical implications. Crit Rev Oncog 2012;17: 1–16.

7. Liu H, Gruber CW, Alewood PF, Moller A, Muttenthaler M. The oxytocin receptor signalling system and breast cancer: a critical review. Oncogene 2020;39: 5917–5932.

8. Kalinina TS, Kononchuk VV, Sidorov SV, Obukhova DA, Abdullin GR, Gulyaeva LF. [Oxytocin receptor expression is associated with estrogen receptor status in breast tumors]. Biomed Khim 2021;67: 360–365.

9. Li D, San M, Zhang J, Yang A, Xie W, Chen Y, Lu X, Zhang Y, Zhao M, Feng X, Zheng Y. Oxytocin receptor induces mammary tumorigenesis through prolactin/p-STAT5 pathway. Cell Death Dis 2021;12: 588.

10. Negro A, Brar BK, Gu Y, Peterson KL, Vale W, Lee KF. erbB2 is required for G protein-coupled receptor signaling in the heart. Proc Natl Acad Sci U S A 2006;103: 15889–15893.

11. Blasco-Benito S, Moreno E, Seijo-Vila M, Tundidor I, Andradas C, Caffarel MM, Caro-Villalobos M, Uriguen L, Diez-Alarcia R, Moreno-Bueno G, Hernandez L, Manso L, et al. Therapeutic targeting of HER2-CB2R heteromers in HER2-positive breast cancer. Proc Natl Acad Sci U S A 2019;116: 3863–3872.

12. Wrzal PK, Devost D, Petrin D, Goupil E, Iorio-Morin C, Laporte SA, Zingg HH, Hebert TE. Allosteric interactions between the oxytocin receptor and the beta2-adrenergic receptor in the modulation of ERK1/2 activation are mediated by heterodimerization. Cell Signal 2012;24: 342–350.

13. Romero-Fernandez W, Borroto-Escuela DO, Agnati LF, Fuxe K. Evidence for the existence of dopamine D2-oxytocin receptor heteromers in the ventral and dorsal striatum with facilitatory receptor-receptor interactions. Mol Psychiatry 2013;18: 849–850.

14. Terrillon S, Durroux T, Mouillac B, Breit A, Ayoub MA, Taulan M, Jockers R, Barberis C, Bouvier M. Oxytocin and vasopressin V1a and V2 receptors form constitutive homo- and heterodimers during biosynthesis. Mol Endocrinol 2003;17: 677–691.

15. Gyorffy B, Lanczky A, Eklund AC, Denkert C, Budczies J, Li Q, Szallasi Z. An online survival analysis tool to rapidly assess the effect of 22,277 genes on breast cancer prognosis using microarray data of 1,809 patients. Breast Cancer Res Treat 2010;123: 725–731.

16. Mihaly Z, Gyorffy B. Improving Pathological Assessment of Breast Cancer by Employing Array-Based Transcriptome Analysis. Microarrays (Basel) 2013;2: 228–242.

17. Cassoni P, Marrocco T, Sapino A, Allia E, Bussolati G. Oxytocin synthesis within the normal and neoplastic breast: first evidence of a local peptide source. Int J Oncol 2006;28: 1263–1268.

18. Zagouri F, Sergentanis TN, Chrysikos D, Zografos CG, Filipits M, Bartsch R, Dimopoulos MA, Psaltopoulou T. Pertuzumab in breast cancer: a systematic review. Clin Breast Cancer 2013;13: 315–324.

19. Belcheva MM, Coscia CJ. Diversity of G protein-coupled receptor signaling pathways to ERK/MAP kinase. Neurosignals 2002;11: 34–44.

20. Hackel PO, Zwick E, Prenzel N, Ullrich A. Epidermal growth factor receptors: critical mediators of multiple receptor pathways. Curr Opin Cell Biol 1999;11: 184–189.

21. Kose M. GPCRs and EGFR - Cross-talk of membrane receptors in cancer. Bioorg Med Chem Lett 2017;27: 3611–3620.

22. Natarajan K, Berk BC. Crosstalk coregulation mechanisms of G protein-coupled receptors and receptor tyrosine kinases. Methods Mol Biol 2006;332: 51–77.

23. Rimoldi V, Reversi A, Taverna E, Rosa P, Francolini M, Cassoni P, Parenti M, Chini B. Oxytocin receptor elicits different EGFR/MAPK activation patterns depending on its localization in caveolin-1 enriched domains. Oncogene 2003;22: 6054–6060.

24. Pequeux C, Keegan BP, Hagelstein MT, Geenen V, Legros JJ, North WG. Oxytocin- and vasopressin-induced growth of human small-cell lung cancer is mediated by the mitogen-activated protein kinase pathway. Endocr Relat Cancer 2004;11: 871–885.

25. Zhong M, Yang M, Sanborn BM. Extracellular signal-regulated kinase 1/2 activation by myometrial oxytocin receptor involves Galpha(q)Gbetagamma and epidermal growth factor receptor tyrosine kinase activation. Endocrinology 2003;144: 2947–2956.

26. Imanieh MH, Bagheri F, Alizadeh AM, Ashkani-Esfahani S. Oxytocin has therapeutic effects on cancer, a hypothesis. Eur J Pharmacol 2014;741: 112–123.

27. Collaborative Group on Hormonal Factors in Breast C. Breast cancer and breastfeeding: collaborative reanalysis of individual data from 47 epidemiological studies in 30 countries, including 50302 women with breast cancer and 96973 women without the disease. Lancet 2002;360: 187–195.

28. Liu H, Muttenthaler M. High Oxytocin Receptor Expression Linked to Increased Cell Migration and Reduced Survival in Patients with Triple-Negative Breast Cancer. Biomedicines 2022;10: 1595.

29. Tchou J, Kossenkov AV, Chang L, Satija C, Herlyn M, Showe LC, Pure E. Human breast cancer associated fibroblasts exhibit subtype specific gene expression profiles. BMC Med Genomics 2012;5: 39.

30. Daub H, Weiss FU, Wallasch C, Ullrich A. Role of transactivation of the EGF receptor in signalling by G-protein-coupled receptors. Nature 1996;379: 557–560.

31. Bertelsen V, Stang E. The Mysterious Ways of ErbB2/HER2 Trafficking. Membranes (Basel) 2014;4: 424–446.

32. Hynes NE, Stern DF. The biology of erbB-2/neu/HER-2 and its role in cancer. Biochim Biophys Acta 1994;1198: 165–184.

33. Moasser MM. The oncogene HER2: its signaling and transforming functions and its role in human cancer pathogenesis. Oncogene 2007;26: 6469–6487.

34. Smith MP, Ayad VJ, Mundell SJ, McArdle CA, Kelly E, Lopez Bernal A. Internalization and desensitization of the oxytocin receptor is inhibited by Dynamin and clathrin mutants in human embryonic kidney 293 cells. Mol Endocrinol 2006;20: 379–388.

35. Conti F, Sertic S, Reversi A, Chini B. Intracellular trafficking of the human oxytocin receptor: evidence of receptor recycling via a Rab4/Rab5 “short cycle”. Am J Physiol Endocrinol Metab 2009;296: E532–542.

36. Jeong J, Kim W, Kim LK, VanHouten J, Wysolmerski JJ. HER2 signaling regulates HER2 localization and membrane retention. PLoS One 2017;12: e0174849.

37. Kirouac DC, D. J, Lahdenranta J, Onsum MD, Nielsen UB, Schoeberl B, McDonagh CF. HER2+ Cancer Cell Dependence on PI3K vs. MAPK Signaling Axes Is Determined by Expression of EGFR, ERBB3 and CDKN1B. PLoS Comput Biol 2016;12: e1004827.

