## Supplementary Material for "Oxytocin receptor and HER2 interactions in breast cancer"

**Table S1 *ERBB2* and *OXTR* mRNA expression information of the 57 cell lines from the Cancer Cell Line Encyclopedia.**

| Breast cancer cell lines | *OXTR* mRNA (log2) | *ERBB2* mRNA (log2) |
|---|---|---|
| CAL120 | 8.16 | 3.57 |
| JIMT1 | 6.93 | 7.61 |
| HS578T | 6.51 | 3.18 |
| HS739T | 6.36 | 3.73 |
| HS281T | 6.29 | 3.76 |
| HS274T | 5.35 | 3.92 |
| HCC1937 | 5.33 | 4.63 |
| BT549 | 4.86 | 3.16 |
| HS742T | 4.45 | 3.14 |
| HS606T | 4.44 | 3.78 |
| HS343T | 3.55 | 3.53 |
| HCC1143 | 2.87 | 4.11 |
| BT20 | 2.62 | 5.11 |
| MDAMB436 | 2.12 | 2.86 |
| CAL851 | 2.07 | 4.69 |
| BT483 | 1.88 | 7.15 |
| MDAMB231 | 1.84 | 3.68 |
| MDAMB468 | 1.71 | 3.58 |
| UACC812 | 1.33 | 10.37 |
| MDAMB361 | 1.17 | 8.62 |
| MDAMB157 | 1.05 | 3.42 |
| CAMA1 | 1.01 | 5.27 |
| HMC18 | 0.97 | 2.80 |
| HDQP1 | 0.90 | 4.83 |
| KPL1 | 0.82 | 4.76 |
| HCC1500 | 0.62 | 4.44 |
| EFM192A | 0.60 | 10.71 |
| MDAMB134VI | 0.51 | 3.47 |
| HMEL | 0.26 | 4.41 |
| MDAMB415 | 0.14 | 5.05 |
| HCC38 | -0.15 | 5.95 |
| MCF7 | -0.20 | 4.65 |
| HCC1599 | -0.33 | 4.48 |
| HCC1395 | -0.35 | 2.97 |
| AU565 | -0.37 | 10.69 |
| CAL51 | -0.41 | 5.25 |
| MDAMB453 | -0.45 | 7.33 |
| ZR751 | -0.49 | 5.75 |

| | | |
|---|---|---|
| BT474 | -0.55 | 9.76 |
| HCC1954 | -0.90 | 10.86 |
| HCC1419 | -0.97 | 10.72 |
| UACC893 | -1.00 | 10.84 |
| HCC1806 | -1.09 | 4.42 |
| HCC2157 | -1.10 | 5.64 |
| **SKBR3** | **-1.11** | **10.10** |
| HCC1187 | -1.58 | 5.71 |
| ZR7530 | -1.66 | 10.57 |
| T47D | -1.70 | 5.54 |
| HCC1569 | -1.70 | 10.08 |
| CAL148 | -1.74 | 6.89 |
| HCC1428 | -1.79 | 4.34 |
| EFM19 | -2.08 | 6.55 |
| DU4475 | -2.44 | 2.18 |
| MDAMB175VII | -2.49 | 6.68 |
| HCC202 | -2.60 | 10.22 |
| HCC2218 | -2.76 | 10.31 |
| HCC70 | -4.73 | 4.37 |

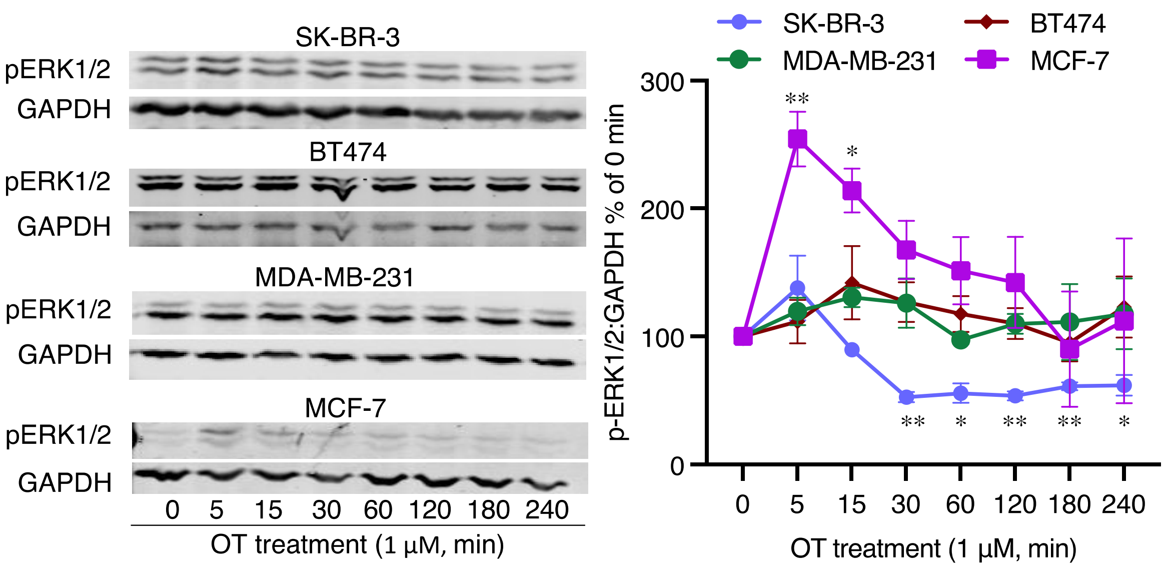


**Figure S1. OT-induced effects on ERK1/2 phosphorylation in different breast cancer cell lines.** Representative Western blot images and quantification of phosphorylated ERK1/2 relative to GAPDH in OT-treated SK-BR-3 (HER2 subtype, $HER2^+$), BT474 (luminal B subtype, $ER^+$ and $HER2^+$), MDA-MB-231 (basal subtype, triple-negative), and MCF-7 (luminal A subtype, $ER^+$) cell lines. *, $p < 0.05$, **, $p < 0.001$ *vs* 0 min group (without OT treatment).
